# How do urban-dwelling bats cope with urbanisation? Sex-dependent plasticity of stress, microbiome and pathogen shedding

**DOI:** 10.1101/2025.05.27.656307

**Authors:** Nia Toshkova, Samantha Aguillon, Magali Turpin, Camille Lebarbenchon, Muriel Dietrich

**Author notes:** **Correspondence** Muriel Dietrich - UMR PIMIT, Sainte-Clotilde, Reunion Island.

## Abstract

Understanding wildlife’s stress and infection response patterns is crucial, especially for species like bats that are ecologically important and potential zoonotic reservoirs. While highly sensitive to habitat destruction and land use change, bats have shown adaptability in human-modified landscapes. This study investigates the impact of urbanisation on the endemic Reunion free-tailed bat (*Mormopterus francoismoutoui*), which roosts in a variety of rapidly urbanised habitats. We analysed 412 urine samples from 11 roosts across Reunion Island, covering a gradient of urban and agricultural land use. We measured urinary cortisol, microbiome diversity and the shedding of *Leptospira* bacteria and paramyxovirus. After accounting for natural cortisol level variations due to circadian rhythm, age, sex, reproduction and capture-induced stress, we found that bats roosting in agricultural and urban areas were in better condition and had lower cortisol levels, suggesting adaptation to human-modified landscapes. Less-stressed bats had more diverse urinary microbiomes and reduced pathogen shedding, suggesting that landscape modifications may indirectly alter the natural epidemiology of bat-borne infectious agents. These patterns were particularly pronounced in females, supporting sex-dependent plasticity to urbanisation in bats. Our study highlights how rapid urbanisation may elicit plastic responses in bats, with potential cascading effects on zoonotic risk of transmission.

## 1. Introduction

Land-use change is one of the most significant anthropogenic disturbances affecting natural ecosystems worldwide and is primarily driven by urbanisation and agricultural expansion [1]. Human encroachment into formerly pristine natural habitats often leads to reduced wildlife populations and a gradual decline in animal biodiversity [2]. Yet, some animals can survive and even thrive in urban environments, exploiting new human-provided resources for foraging and roosting [3]. However, tolerance and adaptation to these human-modified environments may put wildlife under considerable chronic stress [4,5] and deleteriously affect animal health by altering the body condition, immune function and shifting microbiome composition [6–11]. This may thereby increase susceptibility to infections and exacerbate the natural shedding of infectious agents in human-modified landscapes [12,13]. Most of the research on wildlife urban ecology has mainly been conducted on bird species [11,14,15] and thus little is known about the plasticity of stress, microbiome and infection of mammals in response to urban life (but see [16– 19]). Such questions are particularly interesting for mammals involved in zoonoses, as spillovers of emerging infectious agents have been associated with land-use changes and other anthropogenic stressors [20,21].

Bats are the most diverse group of mammals remaining in urban habitats, although they are known to be very sensitive to urban-associated stressors [22,23]. For example, an experimental study showed that gleaning pallid bats (*Antrozous pallidus*) are distracted by noise, which could result in less efficient foraging [24]. Urban-dwelling Kuhl’s pipistrelle (*Pipistrellus kuhlii*) bats also accumulated more metal pollution than their conspecifics in rural areas [25]. Although urbanisation has an overall negative effect on bats, urban tolerance is observed in some bat species and correlates with traits related to greater flexibility in resource requirements and greater mobility [26]. Molossid bats, which are swift aerial insectivorous, are the dominant species within urban bat assemblages. They are frequently found foraging and roosting in urban areas, thus considered well-adapted to human-modified landscapes [27]. However, previous work found that the molossid Brazilian free-tailed bat (*Tadarida brasiliensis*), roosting at human-made bridges in Texas, experienced reduced immune system functioning, which could be linked to physiological stress [28]. This raises the question of whether urban environments genuinely benefit urban-dwelling bats or if bats are merely surviving in these stressful environments [29].

Bats are the natural hosts of several infectious agents and pathogen shedding may be influenced by stress adjustment in response to urbanisation [30]. For example, stress has been linked to (re)activation of viruses and increased viral shedding, such as for Hendra virus and latent herpesviruses [13,31–34]. More recently, high-resolution spatial data showed that anthropogenic stressors (mainly linked to agriculture and deforestation) globally increased coronavirus prevalence in bats [35]. Increased pathogen shedding, in turn, heightens the risk of spillover events, facilitating the transmission of zoonotic diseases to other species, including humans [33]. Thus, understanding the link between land-use modifications, stress, microbiome and pathogen shedding in bats is crucial [30].

In this study, we focused on the Reunion free-tailed bat (*Mormopterus francoismoutoui*), an urban-dwelling molossid bat, to examine the hypothesis of stress adjustment and microbiome change in response to land-use modifications, and its potential consequences for infection. Given the abundance and apparent molossid species’ adaptability to human-modified landscapes, we predict that bats residing in urban and agricultural areas will maintain good health and manage stress effectively, leading to lower susceptibility to infection (prevalence) and reduced pathogen shedding (load). We focused our analysis on the gestation period when females face pregnancy-related physiological stress, which may lead to sex-specific responses to urbanisation [36]. We assessed urinary cortisol levels, bacterial community diversity and prevalence and load of pathogens (paramyxovirus and *Leptospira* bacteria) in bats, across multiple roosts throughout Reunion Island, encompassing a gradient of urbanisation and agricultural levels. We measured and controlled for the natural variation of cortisol levels associated with circadian rhythm, age, sex, body condition and reproductive status of bats, and acute stress induced by the capture and handling of bats.

## 2. Material and methods

### (a) Study sites and landscape characterisation

We studied Reunion free-tailed bats from eleven roosts located all over the island, in either natural settings (caves and cliffs) or anthropogenic habitats (bridges and buildings) (Figure 1). Roost size was estimated visually as described in Aguillon et al. (2023). To characterise the landscape surrounding each roost, land-use data of Reunion Island in 2021 was retrieved from the CIRAD aware website (https://aware.cirad.fr/layers/geonode:OS_2021_SPOT_32740), which includes 28 different habitats. We quantified landscape composition by overlaying a 5-km buffer zone on each roost location, which should encompass the habitat in which bats forage around the roost, based on preliminary tracking data on the hunting distance of Reunion free-tailed bats [37]. The surface of the 28 habitats was obtained for each roost and aggregated into three land-use variables: urban, agricultural and semi-natural (habitats included in each category are listed in Figure S1 and excluded ocean surface, rocks and shadow areas). We used the percentage of each land-use variable in further analyses. To visualise landscape differences among roosts, a Principal Component Analysis (PCA) was applied to the three land-use variables using the *FactoMine* R package [38].

**Figure 1.**
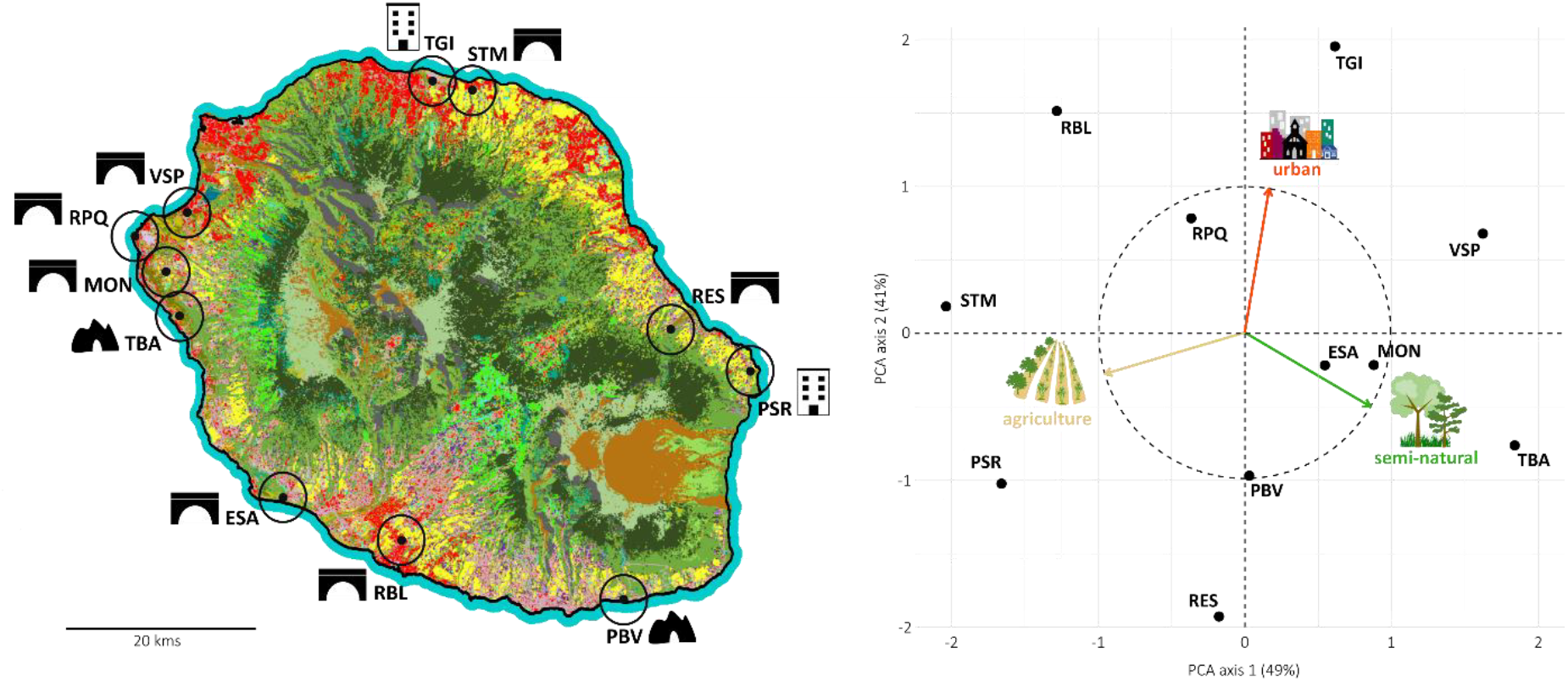
Location of bat roosts and Principal Component Analysis (PCA) across different landscapes in Reunion Island. The roosts are represented by small black points on both the map and the PCA plot. The landscape buffer zone considered for each roost is represented by a black circle on the map and the type of roost (bridge, building, cave or cliff) with a black silhouette. On the PCA plot, the three land-use categories are represented by the arrows.

### (b) Urine sampling

Urine sampling was performed at three different periods of the year: most samples (65%) were collected during a short one-month period in November and December 2021 to avoid temporal variability in cortisol levels. This time of the year corresponds to the gestation period (Aguillon et al., 2023). Additional samples from two roosts, collected in March 2020 when juveniles were present in the population [39], were added to the dataset to investigate the effect of age [40]. Finally, to investigate the circadian rhythm of cortisol levels and measured capture-induced stress, we performed in November 2022, an additional one-time sampling session in a single roost within 12 hours (in the evening and the following morning), using two different methods of urine collection (capture *vs*. trays beneath the roost, see after).

Bats were captured at night after dusk emergence, using harp traps installed close to the roost exit, without preventing the exit of bats. In three roosts (TBA, VSP and STM) where harp trap could not be set up, a butterfly net was used instead by carefully approaching resting (non-flying) individuals during the day (end of afternoon). After capture, bats were immediately hydrated with a sterile syringe and water, placed in a clean individual bag close to a warm source (hot water bottle), and processed at the capture site. A urine droplet was collected while handling bats at the urethral opening, using a pipette and a sterile tip. We also visually determined the sex, age, and reproductive status of each individual. Age was determined by examining the epiphysis fusion in finger articulations that are not welded for juveniles. In females, pregnancy was determined by slight palpation of the abdomen and the presence of inflated nipples. For each bat, we measured the forearm length with a calliper and the mass using an electronic scale to estimate a Body Condition Index (BCI: mass/forearm length ratio). To identify all individuals, bats were tattooed using a dermograph with black ink on the right propatagium with an individual alphanumeric code before being released at the capture site [41]. The time elapsed from capture to release (retention time) was recorded to account for the time span from the onset of handling stress.

To assess cortisol baseline without capturing bats, we collected additional urine samples in the TBA roost (Figure 1) in November 2022, during the evening and the following morning. Urine droplets were collected using sterile plastic film-covered (Saran wrap) cardboard rectangles (20 × 15 cm) placed inside larger plastic trays layered on the ground beneath the roost (as in [42]). Urine droplets are spontaneously shed by flying bats during dusk emergence and in the morning when bats are resting in the colony. All urine samples were stored in sterile vials containing 50ul of Minimum Essential Medium, kept in an ice cooler in the field, prior to transfer to a −80°C freezer.

All fieldwork activities were done using personal protective equipment (FFP2 masks, nitrile gloves, Tyvek suits, and respirator cartridge masks inside caves). During bat capture, gloves were disinfected between each bat individual and changed regularly, and all the equipment was disinfected between sites.

### (c) Measure of urinary cortisol levels

We used the cortisol Express EIA kit (Cayman Chemical Company, USA) and followed the instructions provided by the company to quantify the level of cortisol in urine samples. The assay is based on the competition between cortisol and cortisol-acetylcholinesterase conjugate (cortisol tracer) for the limited number of cortisol-specific mouse monoclonal antibody binding sites precoated on the plate. Urine samples were used at dilution 1:1000 and tested in duplicates. The product of the enzymatic reaction has a distinct yellow colour and was read at 412 nm. Standard curve formulas provided with the kit manual were used to determine the actual concentration of cortisol (ng/mL). Urine samples with %B/Bo values (%Bound/Maximum Bound) greater than 80% (low concentrated) or less than 20% (too concentrated) were re-assayed using dilutions 1:500 and 1:10000, respectively, to fall in the linear range of the standard curve.

We conducted a series of 23 ELISA assays and calculated the intra-and inter-assay coefficients of variation (CV). The intra-assay %CV was 4.2% for the duplicate samples. This is well below the recommended threshold of 10%, indicating high precision in our measurements. Regarding inter-assay CV, which assesses the consistency between different runs or plates, we obtained values of 5.9% and 5.5% for high and low controls, respectively. These values are also below the suggested maximum of 15%, demonstrating good consistency across our assays.

### (d) Shedding of Leptospira bacteria and paramyxovirus

Urine samples were also processed with the Cador Pathogen 96 Qiacube HT kit (Qiagen, Hilden, Germany). Samples were first lysed by mixing 25μL of samples with 175 μL of VXL buffer. Total nucleic acids were extracted in 96-well plates, including negative controls of extraction, and using an automated extractor Qiacube with slight modifications of the Q Protocol, including 350μL of ACB, 100μL of AVE and 30μL of TopElute.

*Leptospira* bacteria and paramyxovirus were detected on a single cDNA preparation. For that, 10 μl of total nucleic acids were submitted to reverse transcription using the Promega cDNA kit (Promega, Madison, USA) with 1.25 μL of Hexamers (Promega), following the manufacturer’s instructions. Five microlitres of cDNA were then used as templates for Polymerase Chain Reactions (PCRs) previously described (Smythe et al., 2002; Tong et al., 2008). Briefly, the presence of *Leptospira* was detected using a probe-specific real-time PCR, targeting a fragment of the 16S rRNA gene. The cycle threshold (CT) of positive samples was noted to infer the relative bacterial load and, thus, the relative intensity of *Leptospira* shedding. For the detection of paramyxoviruses, a semi-nested PCR targeting the polymerase-encoding gene was performed as previously described [43], and no relative viral load could be inferred from this method. Samples collected in March 2020 were already tested in a previous study [40] and thus shedding data were added to the dataset.

### (e) Microbiome sequencing

A subset of extractions of total nucleic acids were used for Illumina sequencing of 16S amplicons (V3-V4 region). Library preparation was performed by GENSOCREEN following the Metabiote® protocol and included negative controls for extraction and library preparation and positive control (mock community provided by GENOSCREEN). Prepared libraries were pooled at equal molarities before sequencing on one lane of an Illumina MiSeq spiked with 15% PhiX using 250 bp paired-end chemistry. We then processed pair-end reads in the FROGS pipeline (“Find Rapidly OTU with Galaxy Solution”; [44]) and clustered them into operational taxonomic units (OTUs) using a maximum aggregation distance of one mutation with the SWARM algorithm [45]. Chimeras were then removed using VSEARCH [46] with de novo UCHIME [47], and a quality filter was carried out by removing any OTU occurring at a frequency below 0.005% in the whole dataset. Taxonomic affiliation of OTUs was performed using NCBI BLAST + automatic affiliation tool available in FROGS pipeline, with the 16S_SILVA_138.1 database. Multi-affiliations were manually checked and resolved with the affiliation Explorer tool on shiny migale (shiny.migale.inrae.fr) when possible. Some OTUs were further collapsed when two conditions were met: OTUs had the same affiliation, and among them, one OTU was always more abundant than the other(s) in several samples. Filtering for potential false positives was done by discarding any count of less than or equal to 5 reads. One OTU exclusively present in the negative control was removed, as well as those (n = 5 OTUs, Table S2) with at least 20 reads in the negative control and present in at least 20% of bat samples.

### (f) Statistical analyses

All statistical analyses and plotting were performed within the R environment, version 4.2.1 [48], using *dplyr* [49], *effects* [50], *ggplot2* [51] and *vegan* [52] packages. Model residuals were inspected with QQplots in the package *DHARMa* [53] and checked for homogeneity of variance, dispersion, and outliers. Then, the ANOVA and chi-square tests were used to test the statistical significance of the explanatory variables in the models.

For stress analyses, cortisol levels were log-transformed to ensure the normality of model residuals. We first investigated the natural variation of cortisol levels over time and the influence of capture by focusing on urine data collected within a single colony (TBA roost) during one day. We fitted a Gaussian generalised linear model (GLM_1_, Table S1), including the timing of collection (evening *vs*. morning) and the method of collection (tray *vs*. capture) as explanatory variables. Then, to assess the variation of cortisol levels, we used a Gaussian GLM on urine data from all captured bats, including individual (age, sex, shedding status), landscape (percentage of urban and agricultural surfaces), and roost (size) factors as explanatory variables (GLM_2_, Table S1). To test if capture-induced stress increased with time, we also included the retention time of captured bats and kept the capture timing in the model. Because of potential sex-specific shedding patterns and responses to landscape modifications, we also included interactions of sex with shedding and landscape variables. Finally, to assess the influence of reproduction on cortisol levels, we used a Gaussian GLM on urine data from captured adult females only (GLM_3_, Table S1), including individual (pregnancy and shedding status) and landscape variables, as well as the retention time and capture timing. To test if stress response of females was modulated differently according to their pregnancy status [36] and urbanisation, we respectively added an interaction in the model between pregnancy and retention time, and between pregnancy and landscape variables.

Variations of the body condition index (BCI) were assessed using a Gaussian GLM from captured adult bats (GLM_4_, Table S1), including individual (cortisol, sex, reproductive and shedding status) and landscape variables, as well as the interactions of sex with shedding and landscape variables. Because BCI was significantly different between adult males and females, higher in females because of the pregnant status of most of them, we repeated this analysis for each sex separately (GLM_4_female_ and GLM_4_male_), and removed the variable ‘reproductive status’ in the male model because males were all in a non-reproductive state at the studied period.

Variations of microbiome diversity were assessed using a Gaussian GLM on urine data from captured adult bats (GLM_5_, Table S1). Alpha-diversity was measured using Hill numbers for *q* = 1, using the R package *iNEXT* [54] and subsequently log-transformed to ensure the normality of model residuals. Explanatory variables included individual (cortisol, sex, reproductive and shedding status) and landscape variables, as well as the interactions of sex with shedding and landscape variables. To analyse the variation of the microbiome at a finer taxonomic resolution, we also used the read counts for the main bacterial family as the response variable in a quasipoisson GLM (GLM_6_, Table S1), including the same explanatory variables as above and the sequencing depth. Finally, to determine whether microbiome composition (beta-diversity) varied with cortisol levels, *Leptospira* and paramyxovirus shedding status, we performed univariate PERMANOVAs on the OTU matrix, using Bray-Curtis distances and the R package *vegan* [52].

To analyse the correlation between cortisol levels and shedding patterns, we first investigated shedding probability using two respective binomial GLMs, one for paramyxovirus (GLM_7_, Table S1) and one for *Leptospira* bacteria (GLM_8_, Table S1). These models were only used with captured adult bats, as too few juveniles were infected (see results). We included individual (sex, reproductive and shedding status, and cortisol level) and landscape variables, as well as the interaction of sex with cortisol and landscape variables. We did not include roost size in the model as a previous study showed no correlation between paramyxovirus/*Leptospira* prevalence and roost size in this bat species [40]. Finally, as *Leptospira* bacteria were detected using a real-time PCR, we used the CT values for PCR-positive adult bats as a proxy for the relative bacterial load in a Gaussian GLM (GLM_9_, Table S1) using the same explanatory variables and interactions as for *Leptospira* shedding probability analysis above.

## 3. Results

### (a) Global cortisol levels and infection prevalence

A total of 412 urine samples were collected and analysed for cortisol levels and *Leptospira* and paramyxovirus shedding status. Among them, 88% (n = 364) were collected while handling captured bats (209 females and 155 males), and 12% (n = 48) were collected from beneath the TBA roost using trays. Cortisol levels ranged from 10.250 to 5756.215 ng/mL. The global shedding prevalence was 40% (± 4.7%) for *Leptospira* bacteria (n = 164 PCR-positives) and 39% (± 4.7%) for paramyxovirus (n = 159 PCR-positives). Co-infections were detected in 21% of samples (n = 88). Shedding was mainly detected in adults, as only three PCR-positives for *Leptospira* bacteria, and two for paramyxovirus, were found among 47 tested juveniles [40]. Prevalence of both infectious agents was higher in adult females compared to adult males (GLM_7_: χ^2^_1_ = 26.215, *P* < 0.001; GLM_8_: χ^2^_1_ = 4.645, *P* = 0.031), and in pregnant females compared to the non-pregnant ones (GLM_7_: χ^2^_1_ = 5.226, *P* = 0.022; GLM_8_: χ^2^_1_ = 10.727, *P* = 0.001).

### (b) Natural variation of stress and effect of handling

The natural circadian rhythm of cortisol showed higher levels in the evening when bats wake up (GLM_1_: χ^2^_1_ = 21.656, *P* < 0.001, Figure 2a). As expected, capturing increased cortisol levels in urine (GLM_1_: χ^2^_1_ = 140.916, *P* < 0.001), especially during the morning. Indeed, following capture, the mean cortisol level was multiplied by 7 times during the evening and 27 times during the morning. Moreover, capture-induced stress increased with retention time of captured bats (GLM_2_: χ^2^_1_ = 20.400, *P* < 0.001, Figure 2b). Roost size was not correlated to cortisol levels (GLM_2_: χ^2^_1_ = 1.361, *P* = 0.188). There was also no difference between sexes (GLM_2_: χ^2^_1_ = 0.025, *P* = 0.857), but we found that adults had more cortisol than juveniles (GLM_2_: χ^2^_1_ = 17.467, *P* < 0.001, Figure 2c). Finally, although cortisol levels were graphically higher in pregnant females in several roosts (Figure S2), we did not find a statistically significant increase in cortisol levels with pregnancy (GLM_3_: χ^2^_1_ = 1.092, *P* = 0.293) and increase in cortisol levels with retention time did not differ between pregnant and non-pregnant females (GLM_3_: χ^2^_1_ = 0.168, *P* = 0.680).

**Figure 2.**
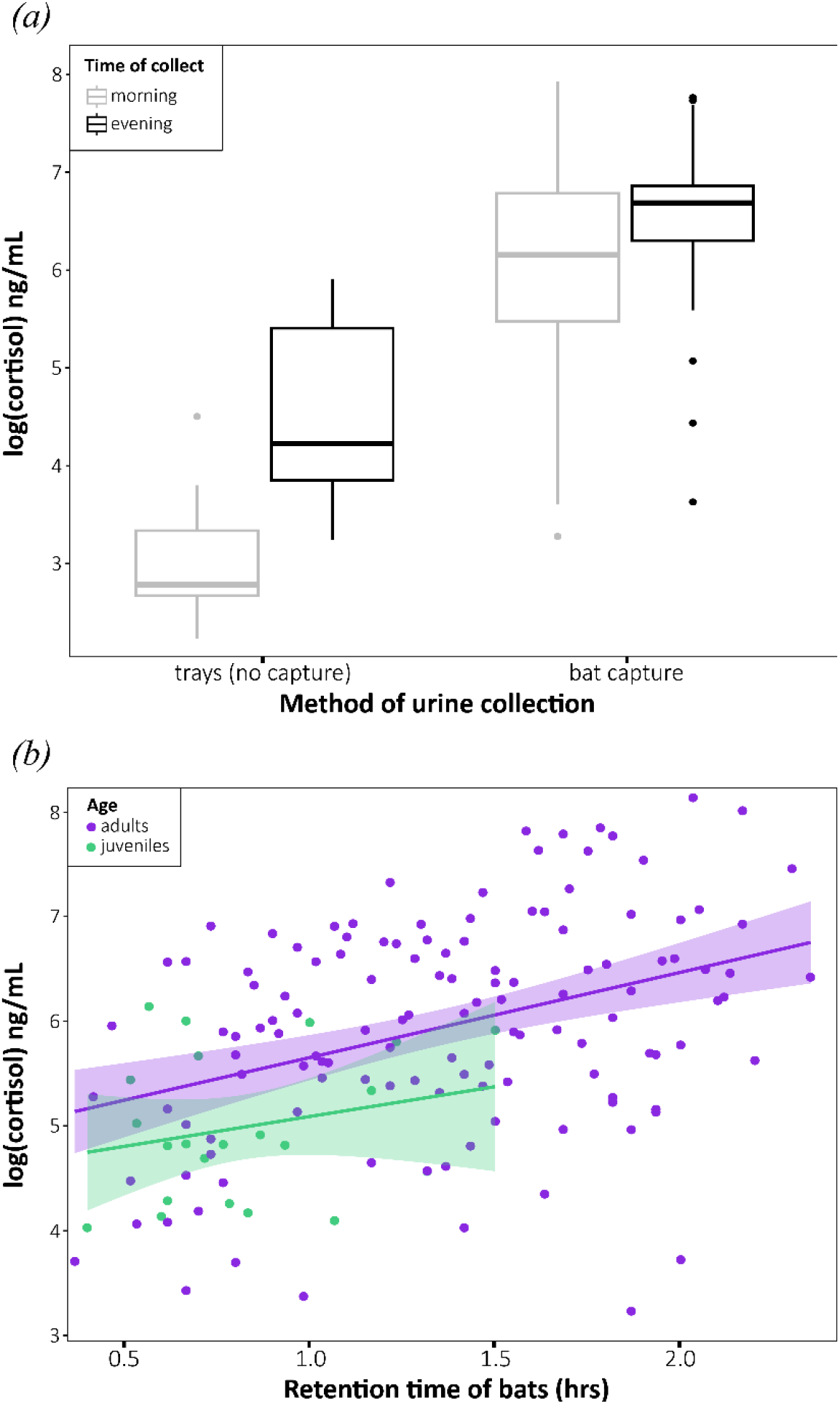
Variation of urinary cortisol levels in Reunion-free tailed bats according to *(a)* timing and method of urine collection, *(b)* retention time and age of individuals. In *(b)*, circles correspond to raw data and the line to predictions of GLM_2_ (see Table S1). Error envelopes depict standard errors below and above the estimated mean responses.

### (c) Influence of landscape surrounding roosts

We found evidence that the landscape surrounding roosts was correlated to the body condition, cortisol levels and infection of bats (Figure 3). Interestingly, when roosting in agricultural areas, bats were in better condition, especially females (GLM_4_: χ^2^_1_ = 0.003, *P* < 0.001; Figure 3a) and had lower cortisol levels (GLM_2_: χ^2^_1_ = 9.096, *P* = 0.001, Figure 3b). Stress response of adult females roosting in agricultural areas was not dependent on their pregnancy status (GLM_3_: χ^2^_1_ = 0.354, *P* = 0.549). In addition, females roosting in agricultural areas were less frequently shedding paramyxovirus (GLM_7_: χ^2^_1_ = 11.965, *P* = 0.001, Figure 3c). This is coherent with the lower probability of paramyxovirus shedding observed in less-stressed bats (GLM_7_: χ^2^_1_ = 5.877, *P* = 0.015), especially in females (Figure 4a), even if there was only a weak evidence of interaction between cortisol and sex in the model explaining paramyxovirus shedding (GLM_7_: χ^2^_1_ = 3.339, *P* = 0.068).

**Figure 3.**
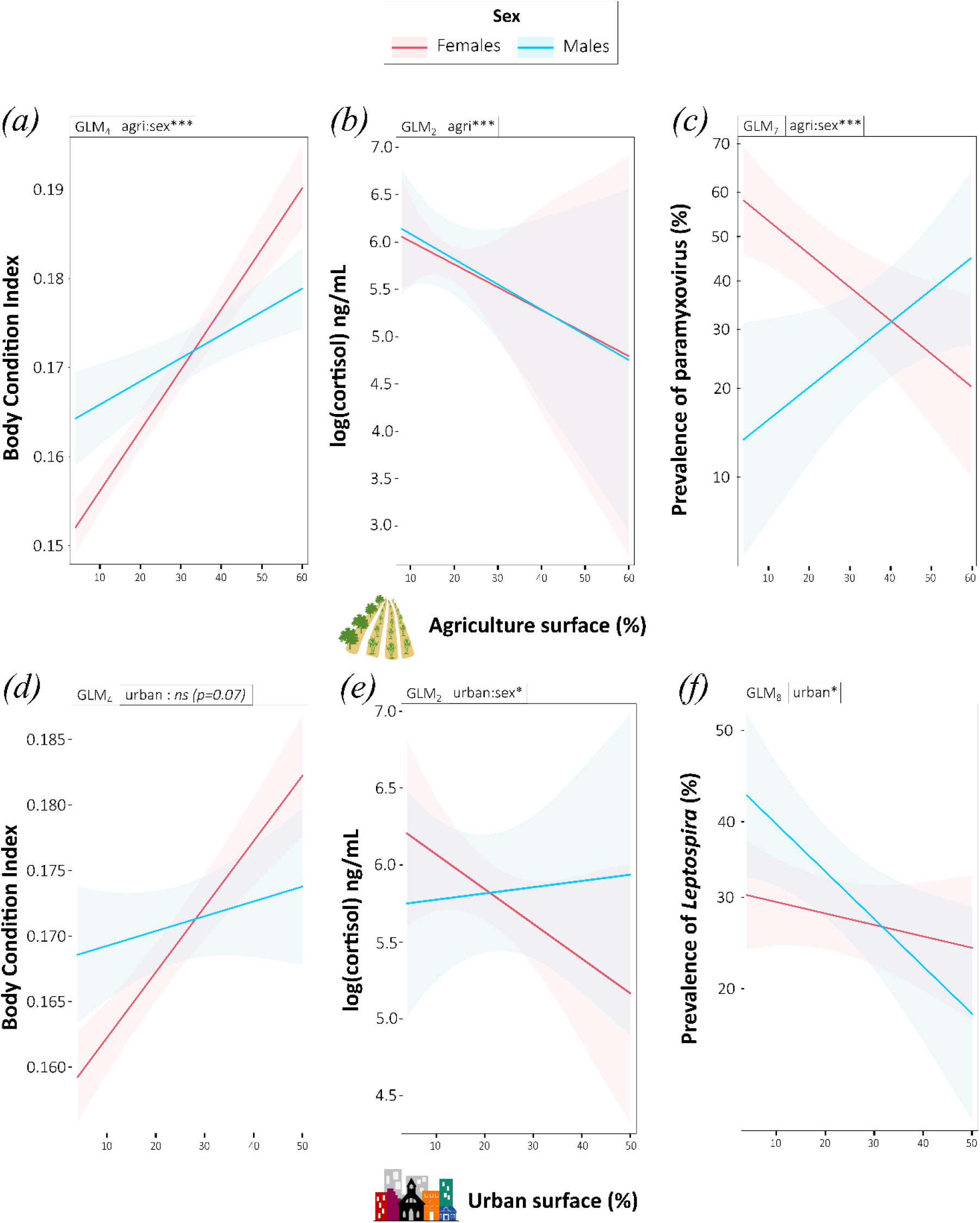
Variation of (a, d) body condition, (b, e) cortisol levels and (c, f) pathogen prevalence in Reunion free-tailed bats in relation to agriculture and urbanisation. Predictions are based on GLMs (see Table S1). The associated GLM is indicated above each graph, with the significance of the plotted variables coded as *ns*: non-significant, *: *p* < 0.05, **: *p* < 0.01, ***: *p* < 0.001. Error envelopes depict standard errors below and above the estimated mean responses.

**Figure 4.**
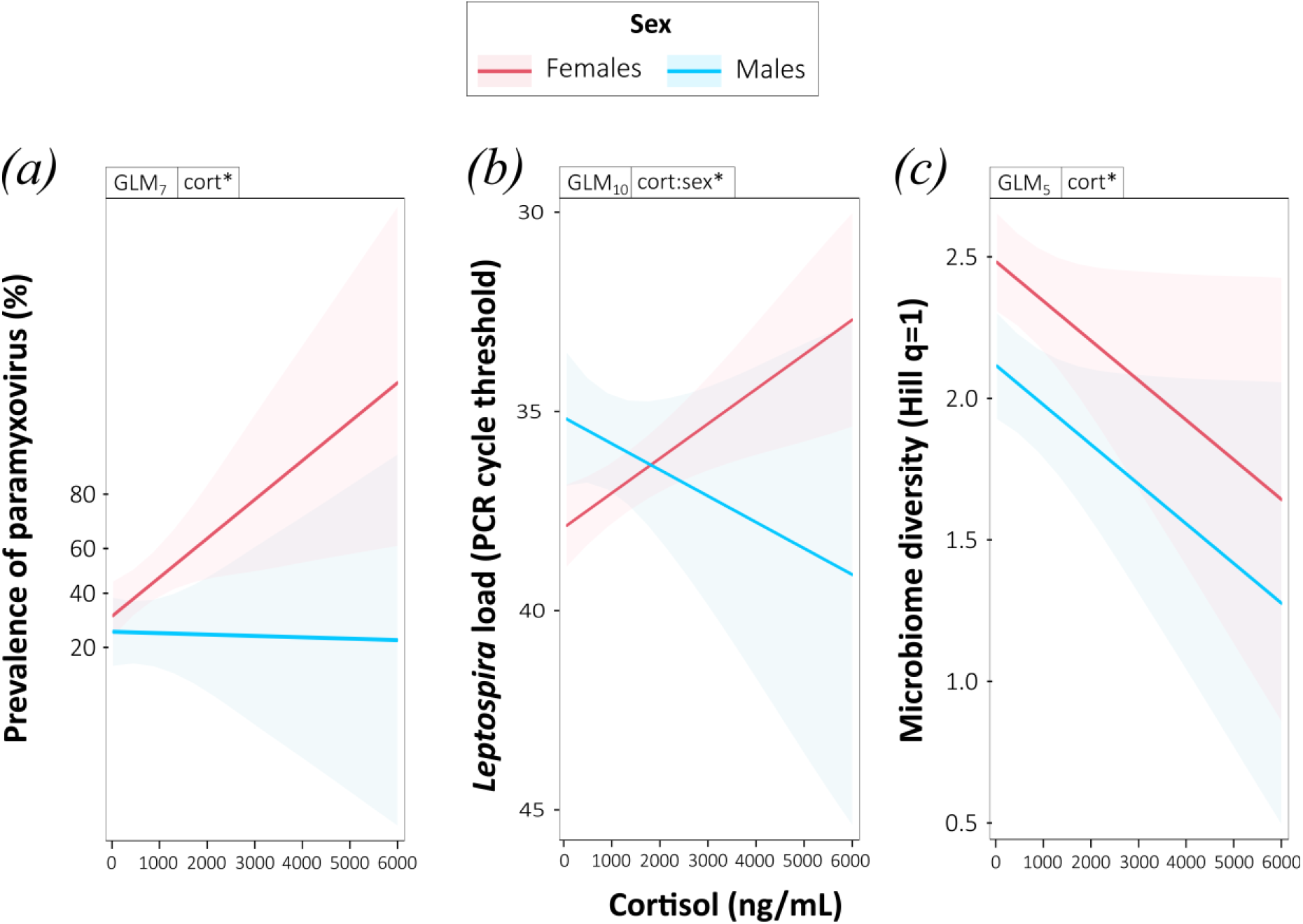
Variation of (a, b) pathogen excretion and (c) microbiome with stress in Reunion free-tailed bats. Predictions are based on GLMs (see Table S1). The associated GLM is indicated above each graph, with the significance of the plotted variables coded as *ns*: non-significant, *: *p* < 0.05, **: *p* < 0.01, ***: *p* < 0.001. Error envelopes depict standard errors below and above the estimated mean responses. In (b), the y-axis is reversed so that lower CT values correspond to higher bacteria load.

In urban areas, only a weak evidence of a similar sex-specific pattern of increased body condition was observed in the model including both males and females (GLM_4_: χ^2^_1_ = 0.000, *P* = 0.077; Figure 3d). When GLMs were performed separately for each sex, body condition was in contrast found significantly higher in urban areas, for both sexes (GLM_4_female_: χ^2^_1_ = 0.002, *P* < 0.001; GLM_4_male_: χ^2^_1_ = 0.001, *P* = 0.037). For cortisol, only females had lower levels in urban areas (GLM_2_: χ^2^_1_ = 4.088, *P* = 0.022, Figure 3e), and this stress response was not correlated to their pregnancy status (GLM_3_: χ^2^_1_ = 0.808, *P* = 0.365). In addition, bats roosting in urban areas were less frequently shedding *Leptospira* (GLM_8_ χ^2^_1_ = 5.418, *P* = 0.020, Figure 3f), but this was not correlated to stress levels (GLM_8_: χ^2^_1_ = 0.004, *P* = 0.949). However, our data showed that *Leptospira* load was reduced when bats had lower cortisol levels, but only in females (GLM_9_: χ^2^_1_ = 39.129, *P* = 0.023, Figure 4b).

### (c) Variation of the urinary microbiome

Finally, we found that females had a higher microbiome diversity than males (GLM_5_: χ^2^_1_ = 3.866, *P* = 0.003) and that the urinary bacterial diversity, but not composition (Bray-Curtis *R*^2^ = 0.003, *P* = 0.739), was negatively correlated to cortisol levels in both sexes (GLM_5_: χ^2^_1_ = 2.218, *P* = 0.024, Figure 4c). Interestingly, the relative abundance of the most prevalent family in the urinary microbiome, Weeksellaceae, increased with increasing cortisol levels (GLM_6_: χ^2^_1_ = 46186, *P* = 0.005). Moreover, urinary bacterial diversity was not affected by *Leptospira* (GLM_5_: χ^2^_1_ = 0.338, *P* = 0.380) nor paramyxovirus (GLM_5_: χ^2^_1_ = 0.007, *P* = 0.897) shedding status, but its composition changed in *Leptospira* PCR-positive samples (Bray-Curtis *R*^2^ = 0.015, *P* = 0.006).

## 4. Discussion

In this study, we used a field-based approach to investigate how human-modified landscapes influence bats and the consequences for the transmission of their infectious agents. We used the endemic Reunion free-tailed bat as a model species of potential adaptation to urban life. To provide a comprehensive view of the overall physiological state of bats, we combined measurements of several biological markers, such as the body condition, baseline and stress-responsive cortisol levels, microbiome diversity and dual infection status. Our findings indicated that bats roosting in agricultural and urban areas effectively managed stress and appeared healthier, exhibiting better body condition, than those roosting in less-disturbed habitats. This supports that urban-dwelling bats, like the Reunion free-tailed bats, are well adapted to living in various human-modified landscapes. Moreover, increased microbiome diversity along with reduced shedding probability and load were found in less-stressed bats, suggesting that landscape modifications by human activities may alter the natural epidemiology of bat-borne infectious agents. These responses were particularly pronounced for females, which also supports sex-dependent plasticity to urbanisation in bats.

### (a) Natural drivers of stress levels

To accurately interpret cortisol levels related to stress, it is crucial to understand the natural factors that influence baseline variations. In this study, we used a dual approach using both capture and no-capture sampling to respectively measure stress-induced and baseline cortisol levels. The cortisol levels in our samples varied widely, ranging from 10.250 to 5756.215 ng/mL. We were able to quantify the influence of capture and retention time and accounted for this bias in the models. We evidenced a dramatic increase in cortisol levels, surging up to 27 times higher when bats were captured during the day. These levels are among the highest ever documented in mammals, surpassing even the previous record values reported from other bat species [55– 57]. This broad range may be due to a tropical syndrome of high stress sensitivity [36], amplified by intense stress response following capture in Reunion free-tailed bats. More broadly, life history traits like long lifespan and low fecundity in bats can also explain higher levels of stress hormones compared to other mammals [58]. Those traits favour higher investment in self-maintenance that might be facilitated by high baseline levels of stress [59].

The validity of our urinary measurements was supported by assessing biologically relevant fluctuations in cortisol levels, such as circadian and age-related variations. In most mammals, cortisol levels follow a circadian rhythm, with peaks occurring just before their active phase [60], and generally rising with age [61]. This age-related increase in cortisol has been demonstrated in several bat species, such as in the Livingstone’s fruit bat (*Pteropus livingstonii*) [62] and the Isabelline serotine bat (*Eptesicus isabellinus*) [63]. Our non-invasive urine collection from beneath the bat colonies revealed that Reunion free-tailed bats exhibit a clear circadian rhythm with higher basal cortisol levels observed at the beginning of the night, which is coherent with their periods of waking and increased activity [64]. The data from captured bats further showed that adults exhibited higher cortisol levels than juveniles, supporting age-related differences in stress response or cortisol metabolism in bats. This may partly result from the ongoing development of the hypothalamic-pituitary adrenal (HPA) axis in early vertebrate life, as well as the possibility that adults develop heightened stress responses over time as a result of their life experiences [65,66].

Group size is known to influence stress responses across animal species [67]. Larger groups can offer benefits such as resource pooling and social support, but also increase stress through parasitism and competition for food, roost space or mates. The wide variation in Reunion free-tailed bat colony sizes, ranging from a few hundred to several dozens of thousands of individuals [39], allowed us to test this effect. However, we did not find such a correlation, consistent with findings in Mexican free-tailed bats [68] and among neotropical bat species [69]. Recent research suggests that the nature of social interactions within a group may often play a more pivotal role in modulating stress responses than the group’s size alone [70]. For example, in captive *P. livingstonii* bats, cortisol levels were linked to the quality of social bonds.[62]. Understanding how the social environment mediates stress response in wild Reunion free-tailed bats could provide valuable insights into how social bonds can buffer against environmental stressors. This knowledge is particularly valuable for conservation strategies, as it suggests that maintaining the quality of social environments may be more effective than focusing solely on population numbers.

Sex-specific stress response have been observed in various bat species, possibly reflecting differences in the HPA axis activity [71], as well as differing ecological demands and behaviours associated with reproduction [36,62,69,72,73]. In pregnant females of a fruit-eating bat (*Artibeus jamaicensis*), Klose et al. [36] found a stronger but shorter acute stress response than in non-reproductive females, likely preventing stress-induced loss of the embryo [36] and supporting the fitness of newborns [74]. However, in our study, no cortisol differences were observed by sex or reproductive status, possibly due to species-specific strategies [62,69] or methodological factors like hormone type or sampling time. For instance, corticosterone, but not cortisol (that we measured here), significantly increased in late-pregnant females [73]. Moreover, our sampling period aligns with the late pregnancy of Reunion free-tailed bats [39]. As stress responses vary across pregnancy stages [61,73], a longitudinal study including lactation, mating, and additional markers like corticosterone, would provide deeper insight.

### (b) Adaptation to urban life?

Complex relationships between stress and extrinsic or intrinsic factors may explain the contradictory physiological responses of wildlife to urbanisation. Indeed, in a recent meta-analysis, Iglesias et al. [5] found no significant differences in stress hormone levels in wild animals between urban and non-urban environments. However, numerous species-specific studies have reported opposite results. In bats, only a few studies have explored how urbanisation affects stress response and they have yielded mixed findings [68,75,76]. One of them performed in flying foxes indicated that urbanisation is associated with chronic stress and poorer body condition [77]. Conversely, a study on *T. brasiliensis* observed no chronic stress in bats roosting in bridges, compared to those in caves [68]. It is further suggested that roosting in human-made structures may improve the health of Brazilian free-tailed bats. Indeed, newborns at the bridge site were in greater body condition and grew faster, compared to those born in a cave [78]. This is in line with our results of reduced urinary cortisol levels and higher body condition in bats roosting in agricultural and urban areas. Response in bats is thus species-specific and we highlight that molossid bats show high ecological plasticity and generally seem to be successful urban adapters [27].

Because urbanisation is relatively recent on Reunion Island (350 years of human colonization), the plastic response of urban dwellers, like the Reunion free-tailed bats, may not be optimal to these rapid environmental changes. As a result, these responses may not provide any evolutionary advantage and could even be maladaptive [5]. Recent genetic analyses suggest that the expansion of the Reunion free-tailed bat’s population is relatively ancient (55,000 years ago) and thus not linked to recent human-modified landscapes [79]. Therefore, to test the true adaptive nature of the observed plasticity in Reunion free-tailed bats, future research should aim at estimating survival and reproductive success through long-term monitoring of bat populations [80]. Genomic and epigenomic analyses can also aid in uncovering the potential molecular mechanisms underlying the observed plasticity in the bat population [81,82].

As bats spend most of their time roosting, roost microclimate (temperature, humidity and windspeed) may strongly influence their physiology and behaviour [83,84]. Understanding these conditions is key for explaining plasticity in Reunion free-tailed bats and many other molossids [27]. Depending on the season, these bats use caves, crevices in cliffs, buildings and bridges [39]. Human-made structures may offer bats a variety of microclimates, allowing them to choose sites that best suit their microclimatic preferences and social needs and consequently significantly reduce stress and improve body condition [68]. For example, Indiana bats (*Myotis sodalis*) show thermoregulatory strategies ranging from normothermia in warm roosts to heterothermia in cooler ones [85], and microhabitat selection varies according to sex, age, and reproductive status [86–88]. Warm environments would provide optimal conditions for pregnant females and their young, while cooler and more buffered environments would promote daily torpor and energy conservation [89]. In little brown bats, reproductive females would rely heavily on building roosts for the duration of the reproductive season [90]. Such sex-specific microhabitat selection may explain the better body condition and lower stress in females in Reunion free-tailed female bats in human-modified landscapes, although we found no clear link to pregnancy.

Dietary plasticity and specific morphofunctional traits of bats may help bats to adapt to urban environments, by enabling effectively foraging in human-modified landscapes [91]. The high-altitude and rapid flight capabilities of molossid bats allow them to access distant food patches scattered across fragmented urban landscapes [27]. The echolocation signals of Reunion free-tailed bats display remarkable structural and frequency variability [92,93], which may also enable them to exploit a broader dietary niche and confer an advantage when encountering new resources in urban environments. This adaptability could be further enhanced by the low interspecific competition on Reunion Island, where only one other insectivorous bat species exists, the Mauritian tom bat (*Taphozous mauritianus*). Recent metagenomic analyses showed that landscape surrounding roosts was correlated to the dietary composition of Reunion free-tailed bats [94]. Bats roosting in agriculture areas consume more Lepidoptera, while those in urban areas increased their consumption of Blattodea. Reunion free-tailed bats could thus take advantage of more predictable and concentrated novel resources in human-modified landscapes, thus reducing their hunting efforts and improving their body condition and overall health. For example, insectivorous bat activity in Australia has previously been shown to increase with greater insect abundance in agricultural environments [95]. Further studies should thus investigate the links between their diet preference, body condition and stress levels.

### (c) Pathogen shedding in human-modified landscapes

Previous experimental studies on birds showed that stress could directly and indirectly mediate the spread of pathogens, by extending the duration of host infectiousness and increasing viral load [96]. In bats, stress has been associated with the (re)activation of latent infections and increased shedding [97], yet studies that directly examine the interaction between stress and infection in bats remain scarce. This gap is particularly significant, as for example, dietary stress, driven by land-use change, is believed to play a key role in mediating the relationship between land-use change and zoonotic spillover events [34]. Here, we uncovered a pathogen- and sex-specific relationship between stress and infection in Reunion free-tailed bats. Specifically, less-stressed female bats roosting in agricultural areas shed paramyxovirus less frequently in their urine. For *Leptospira*, shedding probability was reduced in urban areas for both sexes, but bacteria load was only reduced in less-stressed females that roosted in more urbanised areas. Unfortunately, we could not assess viral load in this study as a qPCR for the specific paramyxoviruses hosted by our model species [42] is not yet designed. Our findings contrast with previous results on flying foxes, showing only a small positive statistical association between Hendra virus shedding status and urinary cortisol concentration [98,99]. However, our findings support our hypothesis of urban-dwelling bats having a lower probability of shedding (for paramyxovirus) and reduced shedding load (for *Leptospira*) in human-modified landscapes, at least during the studied period.

Before concluding that pathogen pressure is lowered in less-stressed Reunion free-tailed bats roosting in human-modified landscapes, further investigations are required to test if patterns of stress, microbiome and pathogen shedding are consistent throughout the year. We recently showed that *Leptospira* and paramyxovirus are shed year-round, and that bats probably host persistent infections with episodic shedding tied to reproductive cycles [40]. Our findings align with earlier data, confirming that sex and reproduction significantly affect shedding prevalence [40]. This variability suggests that the interplay between stress response and pathogen pressure may differ depending on the sex, time of year, and reproductive status of individual bats. Such differences could impact the dynamics of pathogen transmission and ultimately influence spillover risk. Therefore, understanding these seasonal and physiological fluctuations is crucial for assessing the epidemiological implications of land-use change and for predicting periods of heightened spillover potential.

The microbiome can mediate interactions between stress and infection [100] and play a role in host adaptation to urban environments [10,101]. In this study, we focused on the urinary microbiome, known to be highly diverse in bats [102]. Our findings revealed that microbiome diversity was negatively associated with stress levels, though not directly influenced by landscape. Urbanisation can trigger physiological host stress responses that alter immune function, hormone levels, and overall metabolic processes, indirectly shaping microbiome composition [15,103]. As seen in squirrels, these effects act through host physiology rather than direct environmental exposure [9,17]. We found that Weeksellaceae, the most abundant urinary bacterial family, increased with stress. This bacterial family combines aerobic, nonfermenting, oxidase-positive, and Gram-negative rods commonly found in the water, soil, and environment [104]. It has been associated with fungal disease in amphibians [105], and in humans, some species of Weeksellaceae are known to cause diseases such as sepsis, pneumonia, endocarditis and skin infection (*e*.*g*. [106–108]). Our study raises the question of the biological role of such bacterial taxa in bats and highlights the need for a more functional analysis of the bat microbiome.

## 5. Conclusion

Our study sheds light on the intricate relationship between urbanisation, stress, microbiome and infection in bat populations. Reunion free-tailed bats demonstrate remarkable phenotypic plasticity in human-modified landscapes, where they may exhibit reduced transmission of their natural pathogens. This plasticity may not only help bats cope with the rapid urbanisation occurring on Reunion Island but may also precede and guide evolutionary shifts over time [109– 111]. Understanding these processes is crucial because such adaptive traits may influence both the survival of species and the ecological and epidemiological dynamics in increasingly urbanised environments. Long-term monitoring of urban-dweller species like the Reunion free-tailed bat could serve as models for understanding how wildlife responses to anthropogenic changes can influence the risk of zoonotic spillovers.

## Ethics

Bat capture and manipulation were evaluated by the ethics committee of Reunion Island, approved by the French Ministry of Research (APAFIS#10140-2017030119531267), and conducted under permits delivered by the Direction de l’Environnement, de l’Aménagement et du Logement (DEAL) of Reunion Island (DEAL/SEB/UBIO/2018-09) and by the French National Museum of Natural History (CACCHI2022-89A).

## Data accessibility

Metadata of bat samples can be accessed on Zenodo (10.5281/zenodo.15526021). Supplementary material is available online.

## Declaration of AI use

Our use of AI-assisted technology was limited to assisting in revising the wording of portions of our initial draft for clarity, and to help with coding in R language.

## Author’s contributions

N.T.: conceptualization, formal analysis, investigation, writing-original draft, writing-review and editing; S.A.: investigation, writing-review and editing; M.T.: investigation, methodology, writing-review and editing; C.L.: conceptualization, investigation, methodology, supervision, writing-review and editing; M.D.: conceptualization, formal analysis, funding acquisition, investigation, methodology, project administration, resources, supervision, validation, visualization, writing-original draft, writing-review and editing.

All authors gave final approval for publication and agreed to be held accountable for the work performed therein.

## Acknowledgements

We are grateful to Eco-Med Océan Indien, Biotope, the DEER of Région Réunion (Direction de l’Exploitation et de l’Entretien des Routes), the DRT of Département Réunion (Direction des Routes et des Transports of the Department Reunion), and the City hall of Salazie for their help in identifying and accessing bat roosts. We also thank Arline Rajaonarivelo, Clara Castex, Fiona Baudino, Gildas Le Minter, Grégorie Lebeau, Jéremy Dubrulle, Lise Monti, Rachel Leong and Riana Ramanantsalama for their assistance in the field.

