## Supplementary material for "How do urban-dwelling bats cope with urbanisation? Sex-dependent plasticity of stress, microbiome and pathogen shedding"

---

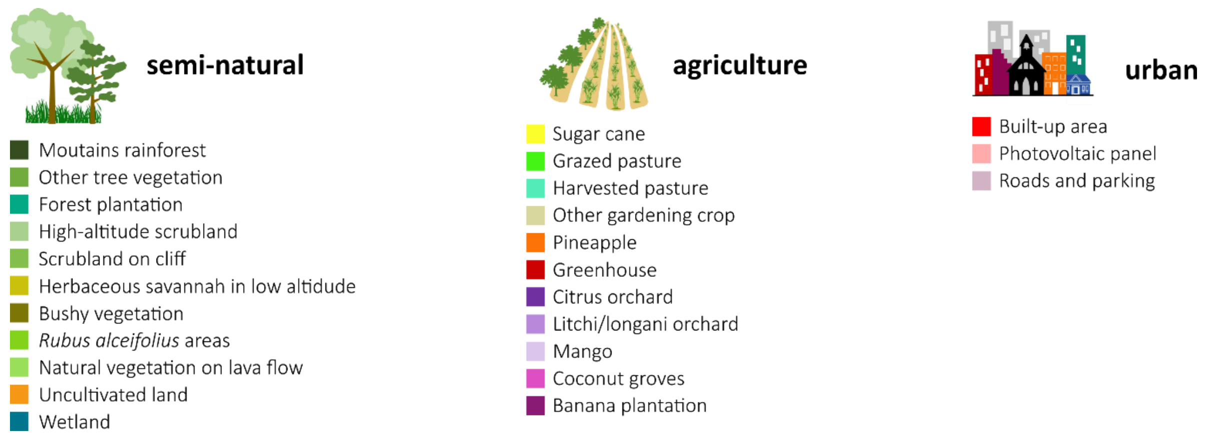


**Figure S1 | List of habitats included in the three land-use variables.** Colors refer to those in the map in Figure 1.

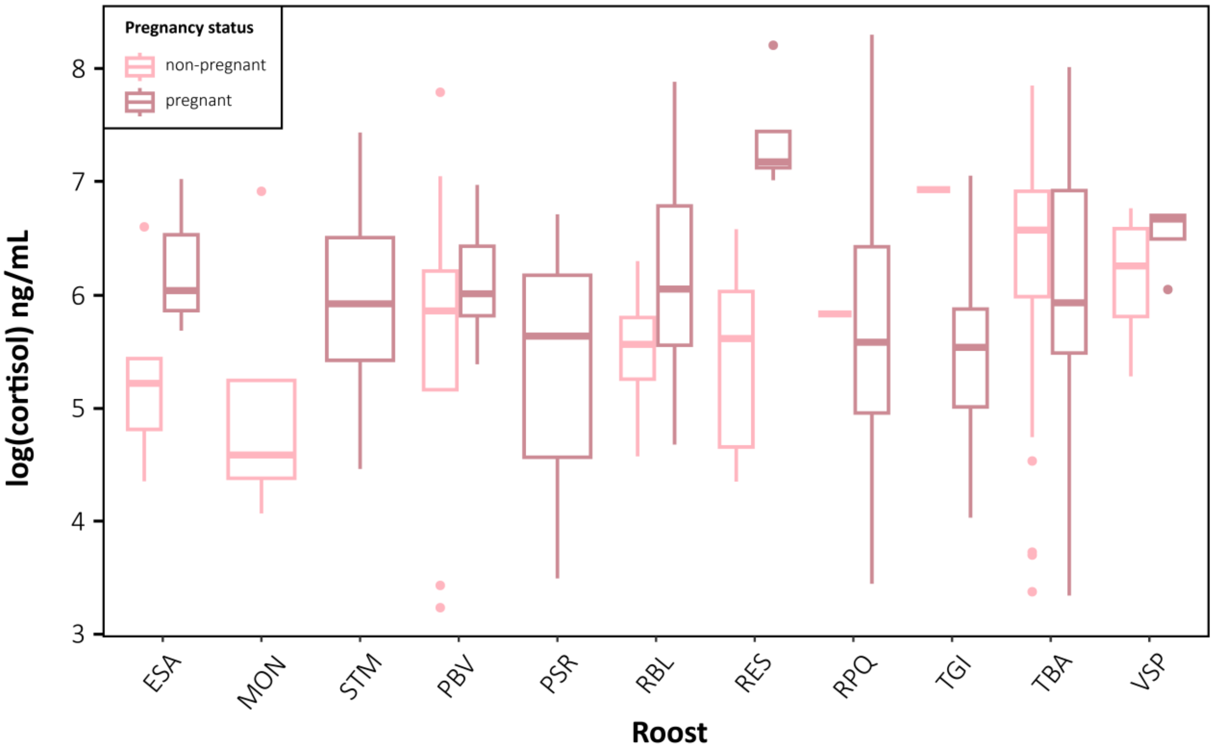


**Figure S2 | Variation of urinary cortisol levels in Reunion free-tailed bats according to the pregnancy status of adult females in several roosts.**

**Table S1 | Summary of the statistical models.** Models include all tested variables and significant variables are underlined and highlighted in bold. N is the number of observations included in the model. The interaction between two variables is represented with a star. Variables are coded as follow: *Timing_collection*: timing of urine collection, *Method_collection*: method of urine collection, *Age*: age of bats, *Sex*: sex of bats, *Urban*: percentage of urban surface around the roost, *Agri*: percentage of agricultural surface around the roost, *Roost_size*: number of bats in the roost, *Lepto*: *Leptospira* shedding in urine, *Pmv*: paramyxovirus shedidng in urine, *Time_manip*: retention time of captured bats, *Preg*: pregnancy status of female bats, *Depth*: sequencing depth of urinary microbiome.

| Variable of interest | Model nb. | Spatial scale of analysis | Origin of urine samples | N | Response variable | Explanatory variables | Df | Deviance | P |
|---|---|---|---|---|---|---|---|---|---|
| Cortisol | $GLM_1$ | Single colony, 1 day | Captured bats and from under the roost | 96 | Cortisol (log) | **Timing_collection** | **1** | **21.656** | **1.185e-06** |
| | | | | | | **Method_collection** | **1** | **140.916** | **< 2.2e-16** |
| | $GLM_2$ | All roosts | Captured bats | 364 | Cortisol (log) | **Age** | **1** | **17.467** | **2.374e-06** |
| | | | | | | Sex | 1 | 0.025 | 0.857 |
| | | | | | | Urban | 1 | 0.706 | 0.343 |
| | | | | | | **Agri** | **1** | **9.096** | **0.001** |
| | | | | | | Roost_size | 1 | 1.361 | 0.188 |
| | | | | | | Lepto | 1 | 0.104 | 0.716 |
| | | | | | | Pmv | 1 | 0.549 | 0.403 |
| | | | | | | **Time_manip** | **1** | **20.400** | **3.407e-07** |
| | | | | | | **Timing_collection** | **2** | **5.657** | **0.027** |
| | | | | | | **Urban*Sex** | **1** | **4.088** | **0.022** |
| | | | | | | Agri*Sex | 1 | 0.635 | 0.368 |
| | | | | | | Lepto*Sex | 1 | 0.434 | 0.457 |
| | | | | | | **Pmv*Sex** | **1** | **2.418** | **0.079** |
| | $GLM_3$ | | Captured adult females | 182 | Cortisol (log) | Preg | 1 | 1.092 | 0.293 |
| | | | | | | **Urban** | **1** | **4.011** | **0.044** |
| | | | | | | **Agri** | **1** | **6.333** | **0.011** |
| | | | | | | Lepto | 1 | 0.107 | 0.741 |
| | | | | | | Pmv | 1 | 2.275 | 0.129 |
| | | | | | | **Time_manip** | **1** | **14.500** | **1.255 e-04** |
| | | | | | | **Timing_collection** | **2** | **6.122** | **0.045** |
| | | | | | | Urban*Preg | 1 | 0.808 | 0.365 |
| | | | | | | Agri*Preg | 1 | 0.354 | 0.549 |
| | | | | | | Time_manip*Preg | 1 | 0.168 | 0.680 |
| Body condition | $GLM_4$ | All roosts | Captured adult individuals | 317 | Body condition index | Cortisol | 1 | 0.000 | 0.825 |
| | | | | | | **Sex** | **1** | **0.003** | **7.920e-06** |
| | | | | | | **Repro** | **1** | **0.027** | **< 2.2e-16** |
| | | | | | | Urban | 1 | 0.000 | 0.077 |
| | | | | | | **Agri** | **1** | **0.024** | **< 2.2e-16** |
| | | | | | | Lepto | 1 | 0.001 | 0.059 |
| | | | | | | **Pmv** | **1** | **0.001** | **0.01160** |
| | | | | | | Urban*Sex | 1 | 0.000 | 0.387 |
| | | | | | | **Agri*Sex** | **1** | **0.003** | **5.926e-06** |
| | | | | | | Lepto*Sex | 1 | 0.000 | 0.996 |
| | | | | | | Pmv*Sex | 1 | 0.000 | 0.981 |

| | | | | | | | | | |
|---|---|---|---|---|---|---|---|---|---|
| | $GLM_{4\_female}$ | | Captured adult individuals - Females | 182 | Body condition index | Cortisol | 1 | 0.000 | 0.255 |
| | | | | | | **Repro** | **1** | **0.027** | **< 2.2e-16** |
| | | | | | | **Urban** | **1** | **0.002** | **1.55e-04** |
| | | | | | | **Agri** | **1** | **0.021** | **< 2.2e-16** |
| | | | | | | Lepto | 1 | 0.000 | 0.204 |
| | | | | | | **Pmv** | **1** | **0.001** | **0.008721** |
| | $GLM_{4\_male}$ | | Captured adult individuals - Males | 135 | Body condition index | Cortisol | 1 | 0.000 | 0.685 |
| | | | | | | **Urban** | **1** | **0.001** | **0.037** |
| | | | | | | **Agri** | **1** | **0.002** | **4.76e-05** |
| | | | | | | Lepto | 1 | 0.000 | 0.18548 |
| | | | | | | **Pmv** | **1** | **0.001** | **0.034** |
| **Microbiome** | $GLM_5$ | All roosts | Adults with paired data (both cortisol and microbiome) | 236 | Bacterial alpha-diversity (log q1) | **Cortisol** | **1** | **2.218** | **0.024** |
| | | | | | | **Sex** | **1** | **3.866** | **0.003** |
| | | | | | | Repro | 1 | 0.157 | 0.547 |
| | | | | | | Urban | 1 | 0.280 | 0.421 |
| | | | | | | Agri | 1 | 0.505 | 0.280 |
| | | | | | | Lepto | 1 | 0.338 | 0.380 |
| | | | | | | Pmv | 1 | 0.007 | 0.897 |
| | | | | | | Urban*Sex | 1 | 0.964 | 0.136 |
| | | | | | | Agri*Sex | 1 | 0.206 | 0.491 |
| | | | | | | Lepto*Sex | 1 | 0.025 | 0.809 |
| | | | | | | Pmv*Sex | 1 | 0.168 | 0.533 |
| | $GLM_6$ | | | | Weeksellaceae (read counts) | **Cortisol** | **1** | **46186** | **0.005** |
| | | | | | | **Sex** | **1** | **224891** | **3.884e-10** |
| | | | | | | Repro | 1 | 185 | 0.858 |
| | | | | | | Urban | 1 | 3436 | 0.439 |
| | | | | | | Agri | 1 | 18735 | 0.071 |
| | | | | | | Lepto | 1 | 32 | 0.941 |
| | | | | | | Pmv | 1 | 6584 | 0.284 |
| | | | | | | **Depth** | **1** | **228750** | **2.753e-10** |
| | | | | | | Urban*Sex | 1 | 7088 | 0.267 |
| | | | | | | Agri*Sex | 1 | 1523 | 0.607 |
| | | | | | | Lepto*Sex | 1 | 6732 | 0.279 |
| | | | | | | Pmv*Sex | 1 | 0 | 0.997 |
| **Infection** | $GLM_7$ | All roosts | Captured adult individuals | 317 | Paramyxovirus shedding | **Sex** | **1** | **26.215** | **3.054e-07** |
| | | | | | | **Repro** | **1** | **5.226** | **0.022** |
| | | | | | | **Cortisol** | **1** | **5.877** | **0.015** |
| | | | | | | Urban | 1 | 1.543 | 0.214 |
| | | | | | | Agri | 1 | 3.328 | 0.068 |
| | | | | | | Lepto | 1 | 2.139 | 0.144 |
| | | | | | | Urban*Sex | 1 | 0.097 | 0.755 |
| | | | | | | **Agri*Sex** | **1** | **11.965** | **0.001** |
| | | | | | | Cortisol*Sex | 1 | 3.339 | 0.068 |
| | $GLM_8$ | | | | *Leptospira* shedding | **Sex** | **1** | **4.645** | **0.031** |
| | | | | | | **Repro** | **1** | **10.727** | **0.001** |
| | | | | | | Cortisol | 1 | 0.004 | 0.949 |
| | | | | | | **Urban** | **1** | **5.418** | **0.020** |
| | | | | | | Agri | 1 | 2.508 | 0.113 |
| | | | | | | Pmv | 1 | 2.142 | 0.143 |
| | | | | | | Urban*Sex | 1 | 3.189 | 0.074 |
| | | | | | | Agri*Sex | 1 | 0.113 | 0.737 |
| | | | | | | Cortisol*Sex | 1 | 0.001 | 0.983 |
| | $GLM_9$ | | Captured adult *Leptospira*-excreting bats | 127 | *Leptospira* load (CT value) | Sex | 1 | 0.479 | 0.802 |
| | | | | | | Repro | 1 | 11.228 | 0.224 |
| | | | | | | **Cortisol** | **1** | **49.489** | **0.011** |
| | | | | | | Urban | 1 | 0.304 | 0.841 |
| | | | | | | Agri | 1 | 2.714 | 0.550 |
| | | | | | | Pmv | 1 | 2.761 | 0.546 |
| | | | | | | Urban*Sex | 1 | 0.558 | 0.786 |
| | | | | | | Agri*Sex | 1 | 0.741 | 0.755 |
| | | | | | | **Cortisol*Sex** | **1** | **39.129** | **0.023** |

**Table S2 | List of OTUs considered as contaminants in the urinary microbiome.**

| OTU_ID | Taxonomic affiliation | Number of reads in the negative control | Prevalence in the bat samples |
|---|---|---|---|
| Cluster_832 | *Jonesia* | 456 | 0% |
| Cluster_3 | *Delftia* | 19801 | 94% |
| Cluster_24 | *Cutibacterium* | 2751 | 89% |
| Cluster_13 | *Schlegelella* | 3127 | 79% |
| Cluster_56 | *Paracoccus* | 4546 | 24% |
| Cluster_354 | *Schlegelella* | 105 | 61% |
